# A Doubly Stochastic Change Point Detection Algorithm for Noisy Biological Signals

**DOI:** 10.1101/106088

**Authors:** Nathan Gold, Martin G. Frasch, Christoph Herry, Bryan S. Richardson, Xiaogang Wang

**Author notes:** Equal Contributors. **Submitted**: Frontiers in Computational Physiology, *The authors gratefully acknowledge technical support by Dr. Qiming Wang and Dr. Michael Last. We would also like to thank Ms. Patrycja Jankowski and Ms. Dana Gurevich for excellent artwork assistance*.

## Abstract

Experimentally and clinically collected time series data are often contaminated with significant confounding noise, creating short, noisy time series. This noise, due to natural variability and measurement error, poses a challenge to conventional change point detection methods.

We propose a novel and robust statistical method for change point detection for noisy biological time sequences. Our method is a significant improvement over traditional change point detection methods, which only examine a potential anomaly at a single time point. In contrast, our method considers all suspected anomaly points and considers the joint probability distribution of the number of change points and the elapsed time between two consecutive anomalies. We validate our method with three simulated time series, a widely accepted benchmark data set, two geological time series, a data set of ECG recordings, and a physiological data set of heart rate variability measurements of fetal sheep model of human labour, comparing it to three existing methods. Our method demonstrates significantly improved performance over the existing pointwise detection methods.

## 1. Introduction

Various biological and medical settings require constant monitoring, collecting massive volumes of data in time series typically containing confounding noise [BP96, GP83, SM90]. This noise, as well as natural fluctuations in the biological system, create non-stationary time series, known as *piecewise locally stationary time series*, which are difficult to analyze in real time [BP77, CP86, Dow07, NFVD11]. Immensely important in clinical and experimental decision making is the accurate and timely detection of pathological changes in the observed time series as they occur. Statistically, this may be interpreted as a change point detection problem for piecewise locally stationary time series [Ada98, ORVSM01, DLRY06].

The heavy contamination of noise due to measurement error and naturally varying phenomena, however, make the detection of change points challenging, as existing techniques will often observe non-pathological changes, resulting in false-alarms and mistrust of detection techniques [BVdA89, Law94, O'C86] - the so-called *cry-wolf* effect. Extracting meaningful change points from naturally occurring fluctuations and noisy corruptions remains a challenge.

The detection of change points in a non-stationary time series is a well-studied problem, which has produced many techniques [BN93, Ada98, TY06, AM07, LS08]. Some of the first work in change point detection is due to [BN93], where changes were detected by comparing probability distributions of time series samples over past and present intervals. This work was extended by Takeuchi and Yamanishi (method TY) [TY06], who proposed a scoring procedure along with outlier detection, to compare past and present probability distributions in real-time. The scoring method is based on an automatically updating autoregressive model, known as the sequentially discounting autoregressive (SDAR) model. Points with scores above a user-defined threshold are declared as change points.

Non-stationary time series may also be viewed as segments of piecewise locally stationary time series [Ada98]. We follow this spirit in our work for the Delta point method. In [Ada98], the locally stationary segments are broken into small pieces and the distance between power spectra for two adjacent pieces are calculated. A variety of distance measures such as the Kolmogorov-Smirnov distance looking at the distance between cumulative power spectra, and the Cramer-Von Mises distance between power spectra was used in [Ada98]. An extension was proposed in [LS08] where the Kullback-Liebler discrimination information between power spectra is used to identify change points. These power spectra methods are particularly well-suited for time series with periodic structure, as they compare power spectra in the frequency domain. Additionally recent advances in real-time change point detection have been made in the machine learning community with promising results, such as relative density ratio estimation method [LYCS13]. Relative density ratio estimation uses a divergence measure to estimate the divergence between subsequent time series samples’ density ratios.

As the current change point detection methodology we consider operates in a pointwise manner, temporal information of change points is lost, such as how often they can be expected to occur, and if they should occur in quick succession or not. Especially in physiological time series, where temporal information and patterns of change points may be highly relevant to practitioners, a pointwise approach may be ill-suited to these time series containing noise. To rectify this problem, we propose a novel change point detection method to analyse the pattern of change points and their inter-arrival times in a small time window so as to observe additional information that may be missed using a pointwise approach. It is our intention this method will reduce the *cry-wolf* effect [Law94] of declaring all of the detected change points as change points relevant to the user. Our method is designed for univariate, piecewise stationary time series, where we seek to correctly classify change points as false alarms or true changes depending on the domain-specific application. The structure of the possibly detected change points may be a shift in mean, change in variance, or the introduction of a new trend to the time series.

While the existing methods [Ada98, BN93, TY06, LS08, AM07], can determine the location of change points, they are not able to extract meaningful change points in time series with piecewise locally stationary structure that contain minor changes and large noise corruption. Our novel method, termed the *Delta point* method, extends the Bayesian online change point detection method of Adams and MacKay [AM07] and later Turner [Tur11], to allow a meaningful change point to be extracted. Our method uses fixed time intervals to construct the joint distribution of the number of change points per interval, and the average length of time between change points in each interval. We demonstrate the effectiveness of the Delta point method on three simulated time series of our own design inspired by existing literature, the widely used Donoho-Johnstone Benchmark curves [DJ94], nuclear magnetic resonance recordings from well-log measurements [óRFP94], annual lowest water levels of the Nile River [Ber94] an ECG recordings from clinical setting [CKH^+^15], and an experimental physiological data set of recordings of fetal sheep heart rate variability during experiments mimicking human labour [WDR^+^14, RJA^+^13, FMM^+^09]. The fetal sheep data set contains short time series with large amounts of measurement noise, while the ECG dataset consists of short time series, with varying features.

## 2. Methods

In this section, we will describe the underlying change point detection methodology our Delta point method extends, as well as the theoretical background of these methodologies. We begin with a notation section for the reader to refer to, and subsequently describe the Bayesian online change point detection methodology introduced by Adams and MacKay [AM07]. Following this, we describe the Gaussian process time series model of Turner [Tur11], which is used as a predictive model for the Bayesian online change point method, and then introduce our Delta point method. The section concludes with a description of the statistical techniques used to analyse the data sets used for testing and our results.

### Notation

Throughout the remainder of the article, we use the following notation (we omit descriptions in this section). Vectors are denoted in **bold**, *eg.*, **y**. *r*_*t*_ is the run length of the change point detection algorithm. **y**_1:*t*_ is a vector of time series observations from time *s* = 1,…, *s* = *t*, with the component *y*_*t*_ the observation at time *t*. **y**_(*r*)_ denotes the vector of time series observations since the previous change point. *τ ϵ* [1, *t* − 1] is a time index denoting the time since the last change point. *p*(·) represents the probability of an event occurring, with *p*(·, ·) denoting a joint probability, and *p*(·|·) a conditional probability. *f* ∼ *GP* (*µ*, *k*) represents a function drawn from a Gaussian process, with *µ* the mean function, and *k*(*·, ·*) is the covariance function or *kernel* for a Gaussian process. *c* = *c*_[*ti,tj*]_ denotes the number of change points in an interval [*t*_*i*_, *t*_*j*_]. 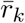 represents the average run length of the change points in the interval [*t*_*i*_, *t*_*j*_]. *α* is a computation parameter, and ℓ is a computation parameter for the input length scale. *ϵ*_*t*_ is Gaussian white-noise, and *σ*^2^ is the standard deviation of the noise. **k**_*\**_ is a vector computed by the covariance matrix, and *K* is a matrix computed by the covariance function.

### Bayesian online change point detection

We begin with a review of the Bayesian online change point detection (BOCPD) algorithm [AM07]. Briefly, BOCPD is a recently introduced change point detection methodology that employs Bayes’ rule to progressively update the probability of observing a change point in time series data while using a predictive model of the time series to make next time step ahead predictions of the time series. The predicted value is then compared with the observed value of the time series to determine if a change-point has occurred.

In detail, BOCPD uses a combination of a predictive model of future observations of the time series, an integer quantity, *r*_*t*_, known as the *run length*, or the time since the last change point, and a hazard function *p*(*r*_*t*_|*r*_*t*−1_), which calculates the probability of a change point occurring with respect to the last change point, to calculate the probability of a change point occurring. Bayes’ rule is used to compute the posterior or past distribution of the run length as

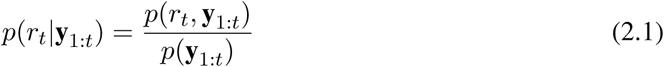

where **y**_1:*t*_ is a vector of past observations of the time series, *p*(*r*_*t*_, **y**_1:*t*_) is the joint likelihood of the cumulative run length and observations, calculated at each step, and *p*(**y**_1:*t*_) is the marginal likelihood of the observations. The joint distribution *p*(*r_t_,* **y**_1:*t*_) is computed with each new observation using a message passing algorithm,

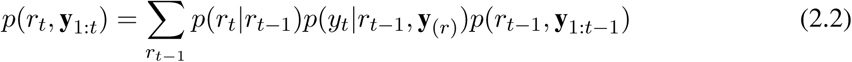
where *p*(*r*_*t*_|*r*_*t*−1_) is the hazard function, *p*(*y*_*t*_|*r*_*t*−1_, **y**_(*r*)_) is the prediction model with observations **y**_(*r*)_ since the last change point, and *p*(*r*_*t*−1_, **y**_1:*t*−1_) is the previous iteration of the algorithm.

### Gaussian process time series

The performance of the BOCPD algorithm is highly dependent on the choice of predictive model for the next time step ahead prediction of the time series data. State of the art performance for the BOCPD algorithm was recently achieved by the use of a Gaussian process time series predictive model [Tur11]. With this in mind, we selected the Gaussian process time series predictive model to be used as the predictive model for the BOCPD algorithm.

A Gaussian process is a Gaussian distribution over functions - that is, the distribution of the possible values of the function follows a multivariate Gaussian distribution [EW06]. Gaussian processes are flexible and expressive priors over functions, allowing patterns and features to be learned from observed data. A Gaussian process is completely specified by a mean function, *µ*(·), and a positive definite covariance function, or *kernel*, *k*(·, ·), which determines the similarity between different observations. The covariance function generates properties of the function drawn from the Gaussian process, such as smoothness and shape. In our work, we used the rational quadratic covariance function,

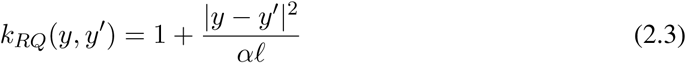

where *α* is a computing parameter, and ℓ is the input scale, or distance between inputs *y* and *y*′, which determines the smoothness of the functions.

In the Gaussian process time series model we used, the time index *t* is used as the input to the function *f* drawn from the Gaussian process, and the time series observation *y*_*t*_ is the output. This is collected in the model,

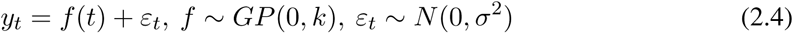

where *ε_t_ ~ N* (0*, σ*^2^) is white-noise in the regression model, and *f ~ GP* (0*, k*) denotes that the function *f* is drawn from a GP with mean 0 and covariance function *k*.

To predict future observations of the time series, we appeal to Bayes’ rule, using the GPTS predictive model as a prior over functions. By the rules of conditioning Gaussian distributions [EW06], the predictive distribution for the next time series observation used in the BOCPD model (2.2) is given by a Gaussian predictive distribution, 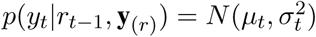, where

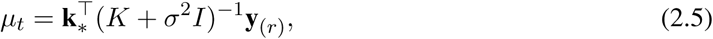

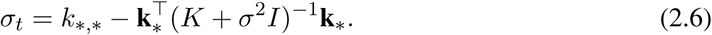

where **k**_*_ = *k*(**y**_(*r*)_, *y*_*t*_) is an (*τ* − 1) *×* 1 vector, where *τ* is the time since the last change-point, *K* = *k*(**y**_(*r*)_, **y**_(*r*)_) is an (*τ* − 1) × (*τ* − 1) matrix, and *k*_*.*_ = *k*(*y*_*t*_, *y*_*t*_).

The GPTS predictive model was then used as the predictive model for the BOCPD algorithm to make next time step ahead predictions of the time series.

### Delta point method

The BOCPD algorithm with the Gaussian process time series predictive model returns a vector of change points. We term the change points stored in this returned vector *suspected change points*. The BOCPD algorithm with the Gaussian process time series predictive model is typically effected by confounding noise in the time series, which in real-world applications is often due to sensor movement or other corrupting sources. Considering the predictive mean and variance from the Gaussian process time series model, equation (2.5) and (2.6), respectively, noise in the observed time series will highly influence the accuracy of predictions. The covariance matrices and vectors in the Gaussian process model are computed on past observations of the time series to be used for forecasting. Highly varied time series observations will thus result in extremely varied future forecast values. The Hazard function in the change point algorithm, *p*(*r*_*t*_ |*r*_*t*−1_), is then updated in the message-passing scheme (2.2) with highly varying information. The probability of change points occurring, as computed by the posterior run length distribution given by equation (2.1), will be set artificially high due to this, thus resulting in over-detection of change points.

The Delta point method is designed to classify, with highest probability, from a vector of suspected change points, the change point most representative of a significant change in the generative process of the time series, and not those given by confounding noise.

The method consists generally of dividing the time series into intervals of user-specified, domain specific length for which a suspected change point may be contained. The number of change points and average run length of the change points in each interval containing a declared change point is then computed, and the interval with the fewest change points and longest average run length is selected as the interval with the highest probability of containing the change point most relevant to the user - the *Delta point*.

Given a vector of suspected change points from the BOCPD algorithm, we can view the observed change points as a realization of a doubly stochastic point process. The doubly stochastic term arises from the intensity function of the change point process being generated from the BOCPD algorithm, which returns a probability of change points occurring. Let {*N* (*t*)}_*t*≥0_ be a counting process representing the number of change points which are declared by the BOCPD algorithm. We assume that {*N*_*t*_}_*t*≥*>*0_ is a doubly stochastic Poisson process, where the intensity is itself a stochastic process, {Λ(*t*)}_*t*≥0_ [DMRT13, Lef07]. Thus, for *t*_2_ ≥ *t*_2_ ≥ 0, we have

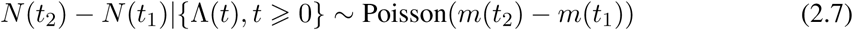

where,

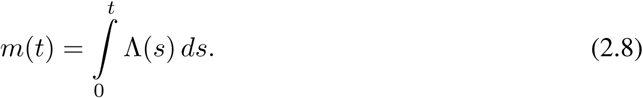

The probability distribution of the intensity function is given by the posterior run length distribution *p*(*r*_*t*_ |**y**_1:*t*_). Since the intensity process is a continuous stochastic process, we need a continuous version of the posterior run length distribution from the BOCPD algorithm (2.2). This is given as

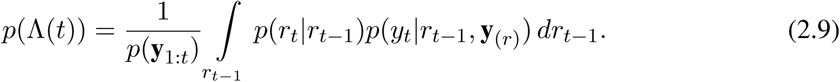

In the time interval (*t*_*i*_, *t*_*i*+1_], where *i* = 1,…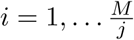 where *M* is the length of the time series, and *j* is the length of each time interval, the probability of observing *k* change points is,

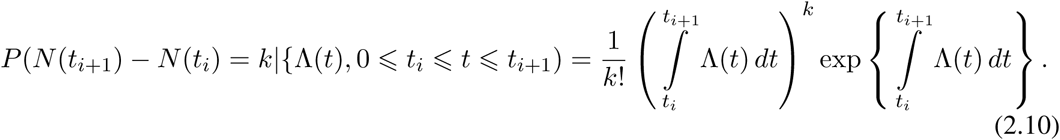

It can be shown that the expected value of the doubly stochastic Poisson process is,

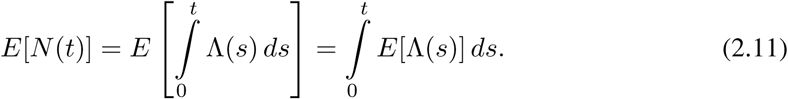

For the Delta point algorithm, the length *j* of the time interval (*t*_*i*_, *t*_*i*+1_] as defined above is user-defined. Let *C*_*i*_ be the number of change points in the interval (*t*_*i*_, *t*_*i*+1_]. Thus, *C*_*i*_ = *N* (*t*_*i*+1_) – *N* (*t*_*i*_), where {*N* (*t*)} is the doubly stochastic Poisson process defined above. The average run length 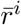 of each interval is computed by the arithmetic mean,

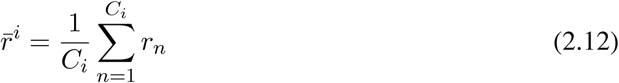

where *r*_*n*_ is the run length associated with each change point in the interval.

Following the average run length computation, we then consider the joint probability distribution of *C*_*i*_ and 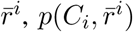, for each interval (*t*_*i*_, *t*_*i*+1_]. The average run length is conditioned by the probability of observing *C*_*i*_ many change-points in the interval

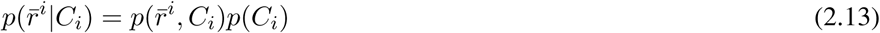

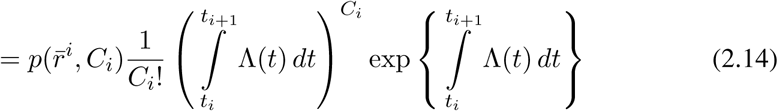

The conditioned averaged run length probability is computed for each interval. Due to the Gaussian process predictive model properties, for a noisy time series, once a change point has been observed in an interval, the probability of more change points being detected in that interval increases. Further, the average run length of that interval decreases accordingly. In the traditional pointwise methods, this will result in many false positives. To avoid this difficulty, we take the opposite approach by observing that the interval with the fewest number of change points and the longest average run length has the highest probability of containing a representative change in the system, and not erroneous change points introduced by noise; that is, it contains a *Delta point*.

Once this interval has been determined, the interval is then searched to look up the associated run length with each interval. The declared change point with the longest average run length *r*_*n*_ is declared as the Delta point.

The only parameter in the Delta point method is the user-defined length of the interval (*t*_*i*_, *t*_*i*+1_], or *j*, as defined above. In this way, the length of the interval generalizes the pointwise change point detection methods, as one can recover the pointwise detection methods by setting the interval length to 1. Conversely, setting the interval length to the length of the time series *M*, will result in selecting the declared change point with the longest run length as the delta point. This follows immediately from taking the arithmetic average of the run lengths. Our method is quite robust to changes in this value, however the best results will be achieved by incorporating expert domain level knowledge to best set the length of the interval, deciding over with time course is of most interest.

In future work, we will explore the structure of the doubly stochastic Poisson process of declared change points in greater detail. As the intensity function is determined predominately by the predictive model, the kernel of the Gaussian process is a natural place to begin investigation. Further, we aim to derive rigorous results on the interval length for optimal performance.

### Statistical analysis

To compare the Delta point method to competing methods, we performed several statistical tests on declared change points from each method. For the simulated data, we computed the mean square error (MSE) for each method, taking the absolute temporal difference between the user labelled change point and the declared change point. We then performed two-sided t-tests to compare the mean absolute detection times of each method to the Delta point method, with the null hypothesis that the mean detection times of other methods will not differ from the Delta point method’s time. For the clinical ECG recording data set, we computed the absolute differences in detection times for each method between the user-labelled change points, and the MSE for each method. We also performed two-sided t-tests to compare the mean absolute detection times between methods with the null hypothesis that the mean detection times of other methods will not differ from the Delta point method’s time. For the experimental data sets, we performed a Fisher’s exact test to compare successful change point detection for each method. In addition, we generated Bland-Altman plots for the fetal sheep physiological data set to compare the accuracy of each method to the user defined change point of interest and the declared change point.

## 3. Results

We tested the Delta point method on several simulated and real world time series data sets. The simulated time series consist of three synthetic time series of our own design, and two widely used benchmark curves. The real world data sets are made up of well-log recordings from geophysical drilling measurements, annual water levels of the Nile river and 100 clinical ECG recordings. We compared the Delta point method to three competing non-stationary change point detection algorithms, namely Takeuchi and Yamanishi (TY) [TY06], Last and Shumway (LS) [LS08], and Liu et al (L) [LYCS13] which were discussed in the Introduction section as popular existing change point detection methods.

### Simulation

To test the efficacy of the Delta point method, we produced 1000 simulations of three different time series, each 500 data points in length. Each time series was designed to simulate change points that may be seen in real world settings, and to have a specific change point that is of more interest than others in the time series. By change point of interest we are are referring to either a change in mean in the case of simulated Series 1 and Series 2, and the introduction of a linear trend in the data in Series 3. These cases are chosen so as to be representative of changes that may occur in real world settings such as sensor failure, or a changing physiological condition.

Series 1 has two change points, with the change point of interest occurring at *t* = 150. This time series simulated the change from a scaled random walk to an autoregressive model, and then back to a scaled random walk. This is a relatively subtle change point to detect, and was inspired from [LS08]. Series 1 is given as,

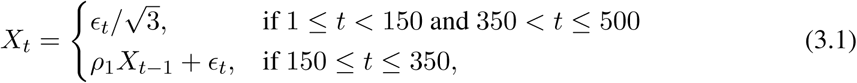

where *ϵ_t_ ~ N* (0, 1), is Gaussian noise, and *ρ*_1_ values are uniformly sampled from [0,1].

Series 2 is a simulated autoregressive model with a large jump, and then a return back to the original process. The change point of interest was chosen as the onset of the jump (*t* = 175). This time series was used to simulate sensor shocks or faults, or a change in the generative parameters of the time series distribution. Series 2 is given as,

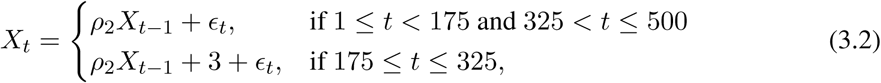

where, *ϵ_t_ ~ N* (0, 1), is Gaussian noise, and *ρ*_2_ values are uniformly sampled from [0,1].

Series 3 is a simulated autoregressive moving average model with an introduced linear trend and subsequent return to the autoregressive moving average model. The change point of interest was chosen as the beginning of the linear trend (*t* = 225). This is a difficult change point to detect, as the noise added to the data obscured the introduction of the trend. This time series simulated the accumulation of some product in a system. Series 3 is given as,

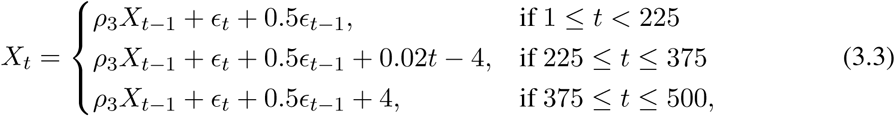

where, *ϵ_t_ ~ N* (0, 1), is Gaussian noise, and *ρ*_3_ values are uniformly sampled from [0,1]. Simulated time series from Series 1 – Series 3 are displayed in Figures 1-3, respectively. The parameter values used for the simulation are *ρ*_1_ = 0.7*, ρ*_2_ = 0.4, and *ρ*_3_ = 0.5.

**Figure 1:**
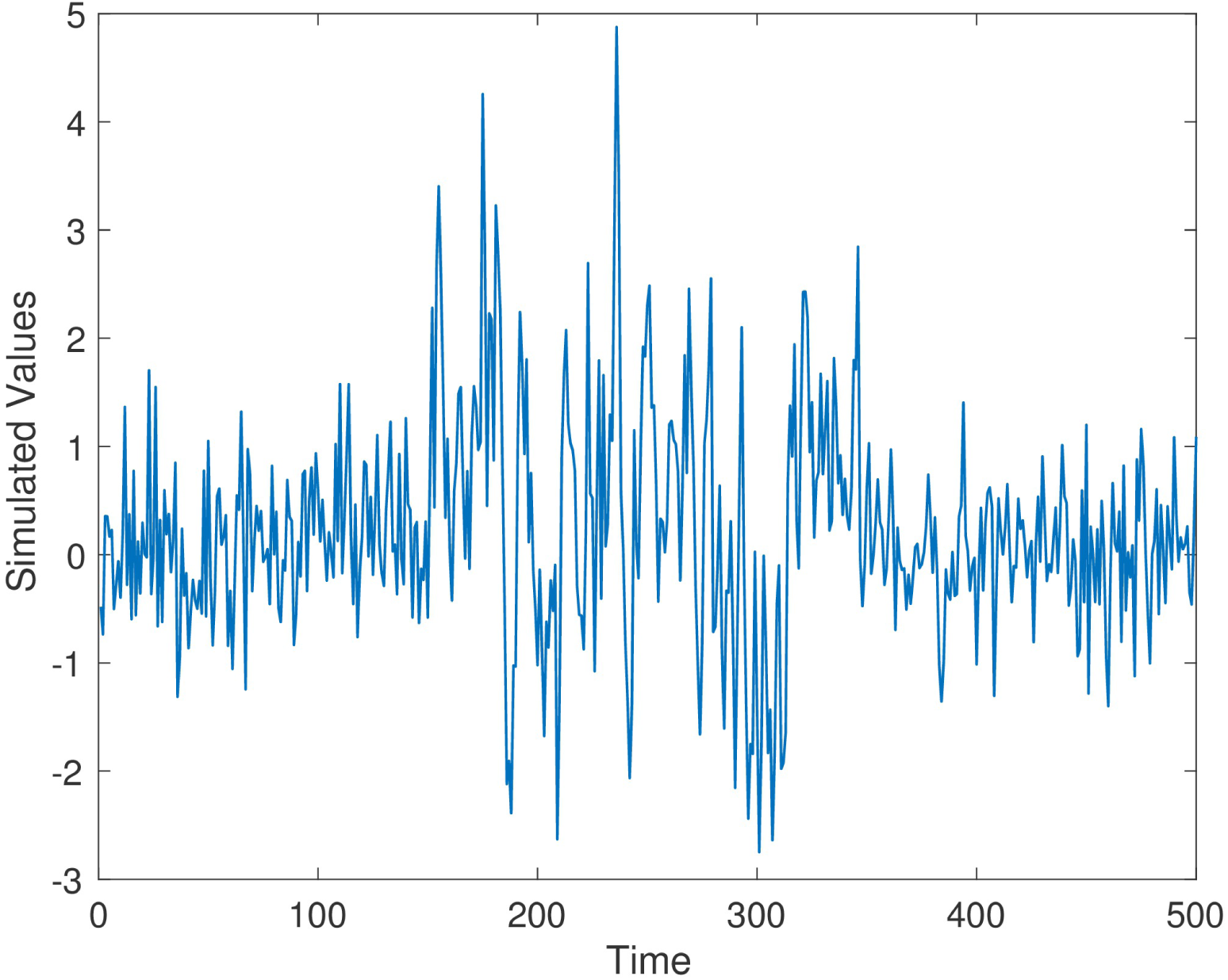
Simulation Time Series 1, *ρ*_1_ = 0.7.

**Figure 2:**
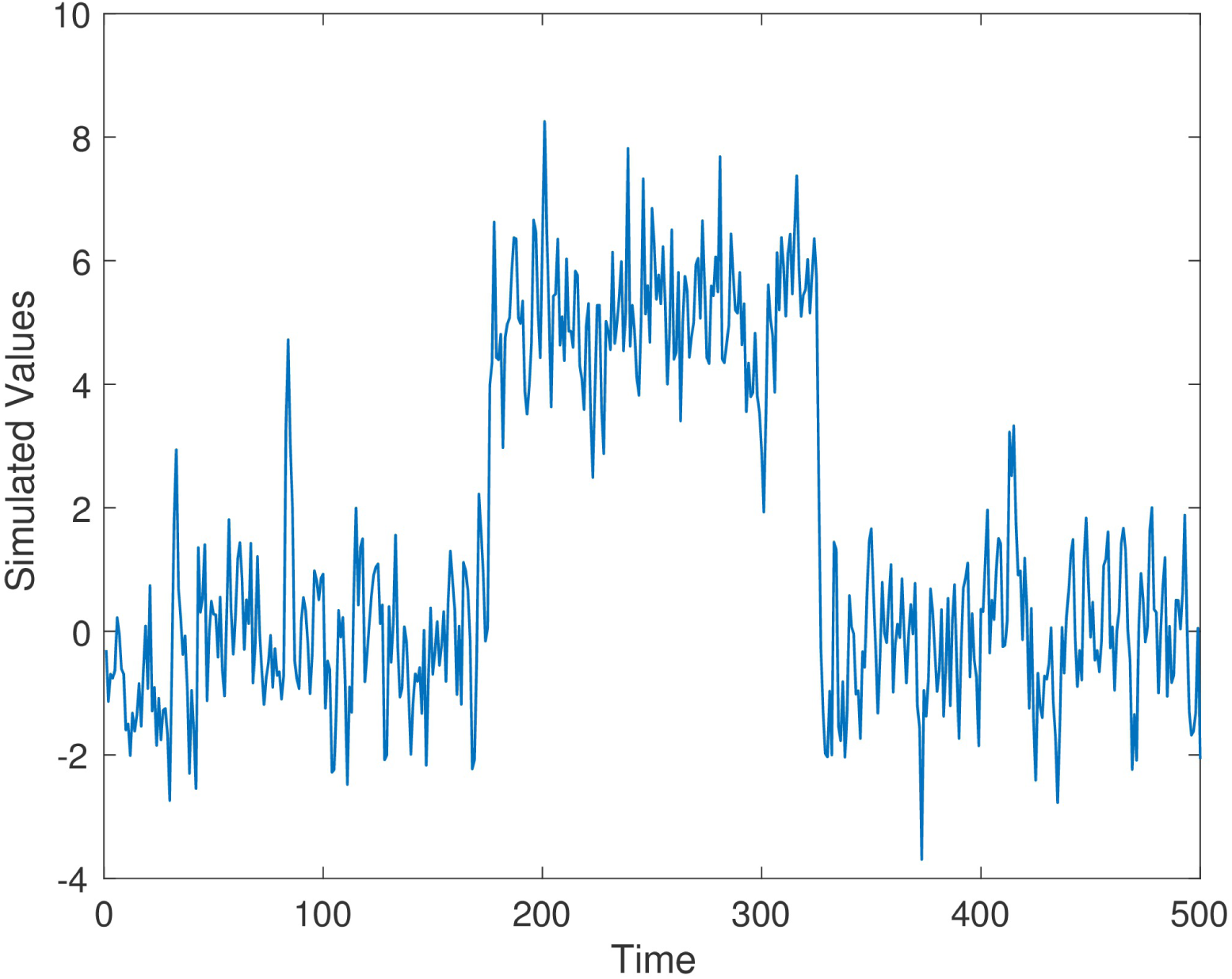
Simulation Time Series 2, *ρ*_2_ = 0.4.

**Figure 3:**
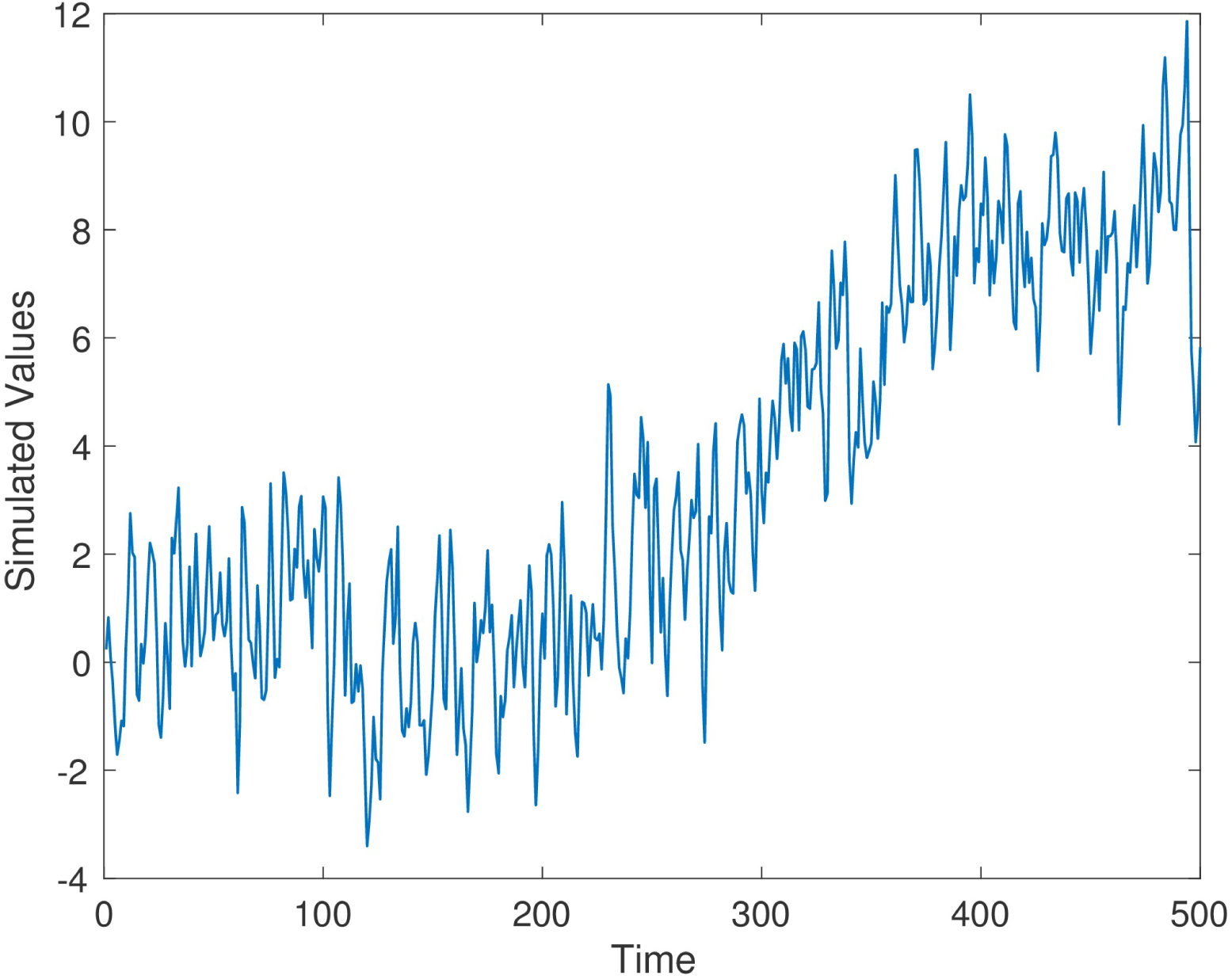
Simulation Time Series 3, *ρ*_3_ = 0.5.

The BOCPD method learned parameter values through training, so we only list the values we used to initiate the method. For Series 1, Series 2, and Series 3, we used a training set of 200 data, taken at the beginning of the time series. The Gaussian process model used a non-biased parameter initialization, with an assumed standard normal distribution prior for Series 1, Series 2, and Series 3. The hazard rate parameter used for the hazard function for initial training for each time Series is *θ*_*h*_ = − 3.982. The Delta point interval length for each time series was set at 40 for training, as this should protect against the BOCPD possibly declaring too many erroneous change points, by being set too short. Setting the interval length to longer should produce similar results. The techniques TY, LS, and L all require a threshold value above which a change point will be declared. We performed cross-validation of several threshold values for each method, choosing the value for each time series that allows the most accurate detection of the change point of interest. We select the change point declared by each method that is closest to the significant change point described above.

The results of each method are displayed in Table 1. We report results for *ρ*_1_ = 0.7*, ρ*_2_ = 0.4, and *ρ*_3_ = 0.5, respectively; the results for different values are *ρ*_1_*, ρ*_2_, and *ρ*_3_ are not significantly different. For Series 1, the Delta point method significantly (*p* < 0.001) outperformed methods TY and LS in the mean absolute difference (absolute difference) of detection time, and had a significantly lower MSE. This difference in performance is confirmed by a two sided t-test with a null hypothesis that other methods do not have a significant mean absolute difference from the Delta point method. In our simulations, method L had a slightly smaller mean absolute difference (8.953) compared to the Delta point method (9.718), however the distribution of declared change points had a larger standard deviation (12.783 compared to 9.881). As well, the Delta point method had a lower MSE. The two sided t-test confirmed that both methods had indistinguishable performance (*p* = 0.135). For Series 2, the Delta point method significantly (*p* < 0.001) outperformed all methods. For Series 3, method TY performed the best, with the lowest mean absolute difference. Figures 4-6 display box plots of the results of each method for Series 1-3, respectively.

**Figure 4:**
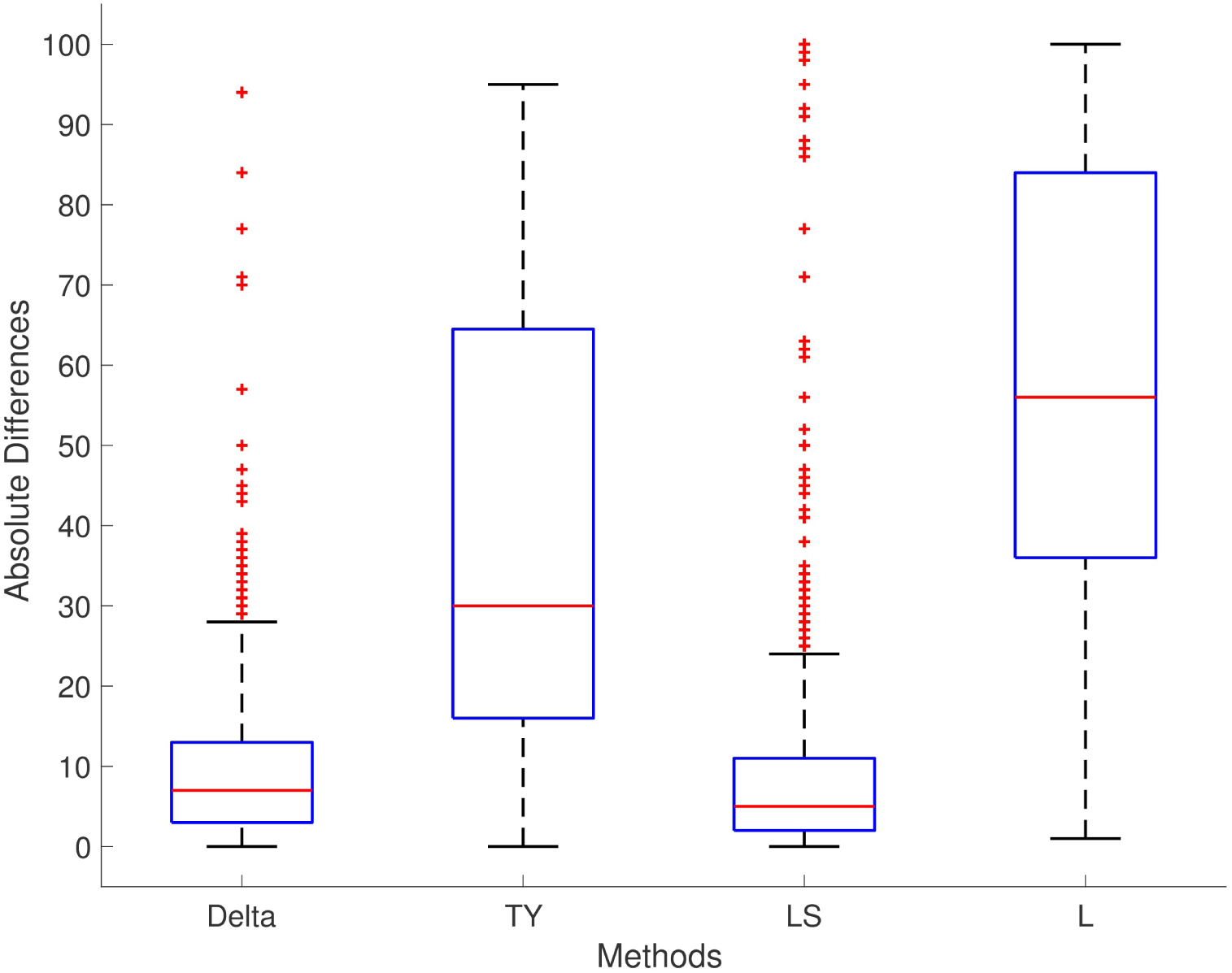
Simulation Series 1 boxplot. Legend: TY = [TY06], LS = [LS08], L = [LYCS13]. Box-plot of absolute differences of detected change points for 1000 simulations of simulation data set Series 1. The true change point location is located at 0.

**Figure 5:**
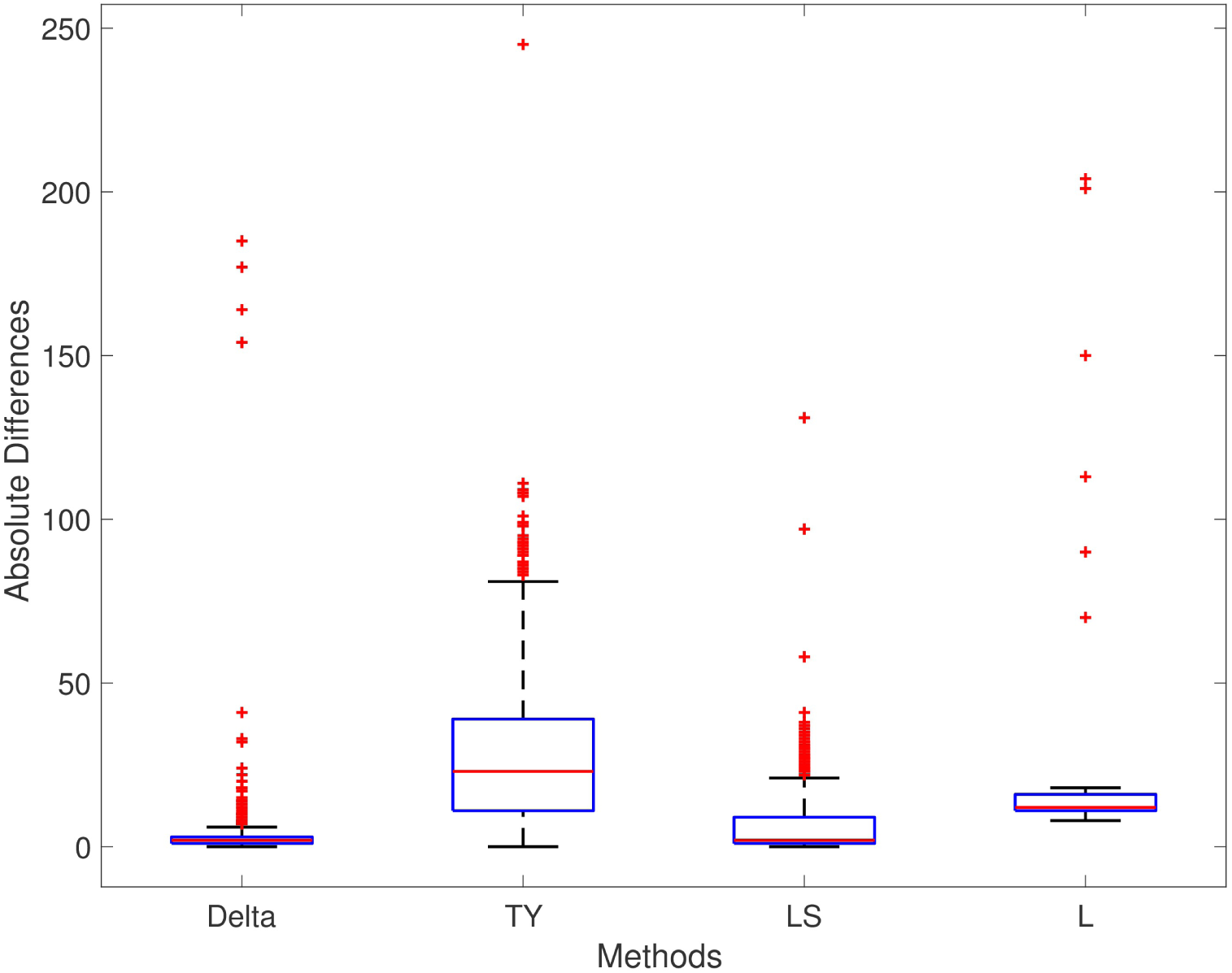
Simulation Series 2 boxplot. Legend: TY = [TY06], LS = [LS08], L = [LYCS13]. Box-plot of absolute differences of detected change points for 1000 simulations of simulation data set Series 2. The true change point location is located at 0.

**Figure 6:**
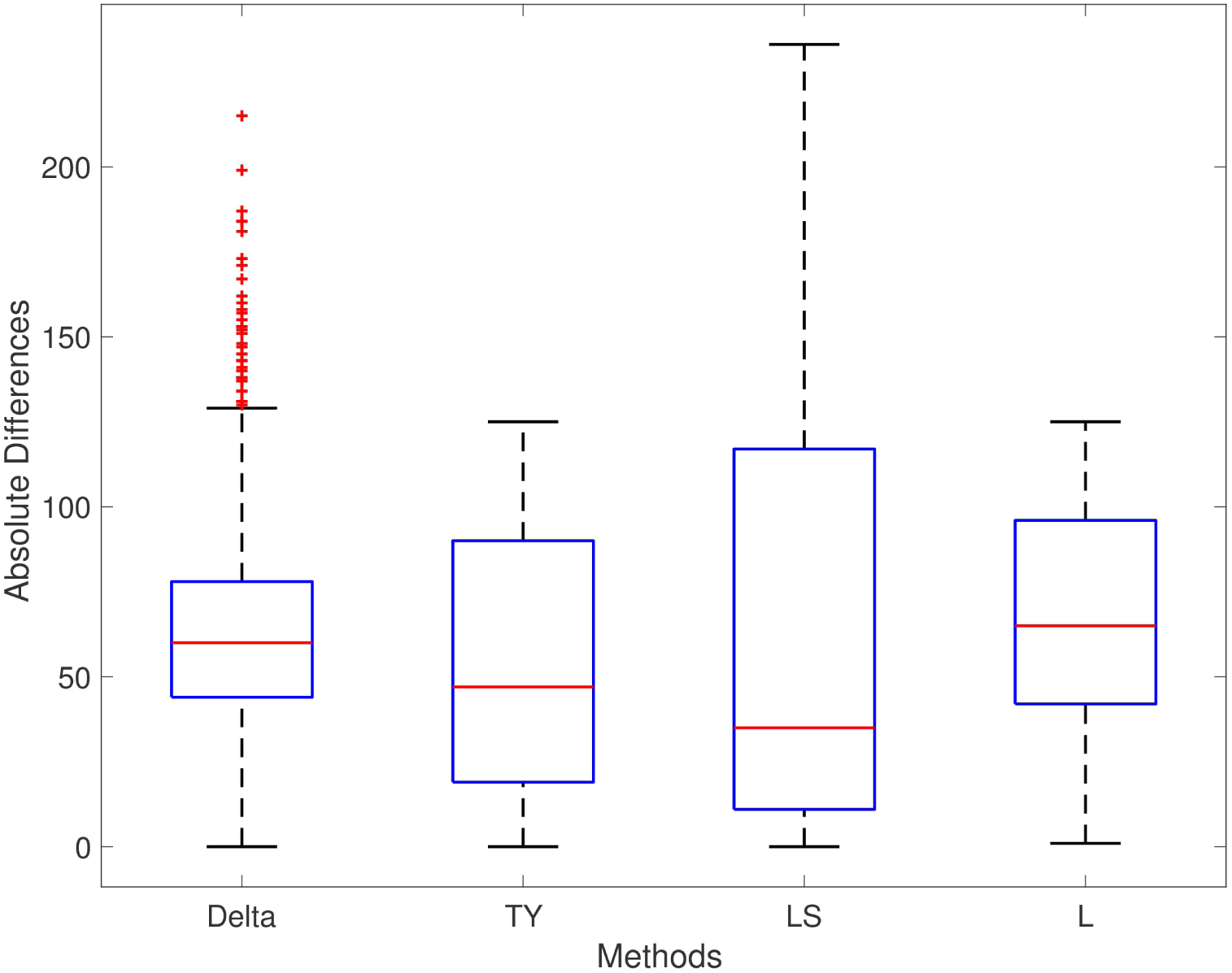
Simulation Series 3 boxplot. Legend: TY = [TY06], LS = [LS08], L = [LYCS13]. Box-plot of absolute differences of detected change points for 1000 simulations of simulation data set Series 3. The true change point location is located at 0.

### Donoho-Johnstone Benchmark

To further analyse the performance of the Delta point method, we tested it and the existing methods on the Donoho-Johnstone Benchmark non-stationary time series [DJ94]. The Donoho-Johnstone Benchmark is a classic collection of four non-stationary time series designed as a test for neural network curve fitting. The curves are known as the Block, Bump, Doppler, and Heavisine, and are 2048 data points in length each. We adapted the curves with the introduction of noise to test for change point detection. As the Delta point method is not designed to function with time varying periodic data, rather piecewise locally stationary time series, we did not test with the Doppler and Heavisine curves.

**Table 1:** Simulation results

| Method | Abs. Diff. (mean ± st. dev.) | MSE $\times 10^3$ | Significance ($p$-value) [CI] |
| --- | --- | --- | --- |
| Series 1 |  |  |  |
| Delta | 9.718 ± 9.881 | 0.192 | N/A |
| TY | 40.110 ± 28.367 | 2.413 | 1 ($p < 0.001$) [-32.255, -28.539] |
| LS | 8.953 ± 12.783 | 0.243 | 0 ($p = 0.135$) [-0.237, 1.767] |
| L | 59.459 ± 25.713 | 4.196 | 1 ($p < 0.001$) [-51.449, -48.033] |
| Series 2 |  |  |  |
| Delta | 3.399 ± 12.099 | 0.158 | N/A |
| TY | 13.802 ± 10.7298 | 0.306 | 1 ($p < 0.001$) [-11.906, -9.406] |
| LS | 6.205 ± 9.6699 | 0.132 | 1 ($p < 0.001$) [-3.7662, -1.8458] |
| L | 27.728 ± 22.961 | 1.296 | 1 ($p < 0.001$) [-25.938, -22.719] |
| Series 3 |  |  |  |
| Delta | 63.603 ± 30.762 | 4.991 | N/A |
| TY | 54.645 ± 39.358 | 4.534 | 1 ($p < 0.001$) [5.860, 12.056] |
| LS | 65.263 ± 63.469 | 8.284 | 0 ($p = 0.457$) [-6.304, 2.714] |
| L | 69.911 ± 30.418 | 5.812 | 1 ($p < 0.001$) [-8.991, -3.625] |

Legend: TY = [TY06], LS = [LS08], L = [LYCS13]. Comparison of methods for detecting significant change point for 1000 simulations of simulated time series: Series 1, 2, 3, respectively; 500 observations in length each. Mean ± standard deviation of absolute difference (Abs. Diff.) between labelled times and detected time of each method are given. Mean Square Error (MSE) of each method is displayed where the lowest value displayed had the least detection error. Results of null hypothesis two-sided t-test comparing absolute differences to Delta point method displayed with *p*-values and confidence intervals [CI].

Legend: TY = [TY06], LS = [LS08], L = [LYCS13]. Comparison of change point detection results for two Donaho-Johnstone Benchmark curves (Bump and Block) with user-labelled change points. User-labelled change points are selected to represent a drastic change in the time series (Bump), or a significant shift in the mean (Block).

For training for the Delta point method, we used a standard normal distribution prior for the Gaussian process, and hazard rate parameter *θ*_*h*_ = − 3.982. We set the Delta point interval to 50 for the Bump curve and the Block curve. We selected 50 time points for the interval length so that we could observe sufficient temporal structure for the doubly stochastic Poisson process. The training set consisted of the first 800 data of each curve, to correspond to the rule of thumb of using the first 35% to 40% of time series data for training [Tur11].

The results of the methods for the Bump and Block curves are displayed in Table 2. The Delta point method performed very well in these cases, declaring the change point of interest very close to the user-labelled point. The Bump curve was a more difficult curve to detect change points in, due to noise. The Delta point method and LS curve have about the same performance for the bump curve (absolute difference of 5 and 4, respectively). The Delta point method outperformed methods TY, LS, and L for accurate detection in the Block curve (absolute difference compared to 17, 22, and 24, respectively).

**Table 2:** Donaho-Johnstone Benchmark curves results

| Method | Detected time | Abs. Diff. |
| --- | --- | --- |
| Bump | Labelled: 440 |  |
| Delta | 445 | 5 |
| TY | 475 | 34 |
| LS | 444 | 4 |
| L | 468 | 28 |
| Block | Labelled: 1331 |  |
| Delta | 1332 | 1 |
| TY | 1348 | 17 |
| LS | 1353 | 22 |
| L | 1335 | 24 |

### Well-log and Nile recordings

The well-log data set consists of 4050 nuclear magnetic resonance measurements obtained during the drilling of a well [ÓRFP94]. These data are used to deduce the geophysical structures of the rock surrounding the well. Variations in the mean reflect differences in stratification of the Earth’s crust [AM07]. The well-log data set is a well studied time series for change point detection [ÓRFP94, AM07, Tur11]. We selected the largest jump in the mean of the time series as the most significant change point (*i* = 1070).

Legend: TY = [TY06], LS = [LS08], L = [LYCS13]. Comparison of detected change point of importance in nuclear magnetic resonance measurements from a rock drill used to detect changes in rock stratification, and lowest annual water levels of the Nile River from 622-1284. The change point of importance is selected as the first significant jump in the mean, indicating the presence of a change in the ground rock, and the instillation of the Nilometer, respectively.

The Nile river time series consists of a record of the lowest annual water levels between 622-1284 CE, recorded on the Island of Roda, near Cairo, Egypt [Ber94]. The Nile river data set has been used extensively in change point detection [Tur11], making it an effective benchmark for the Delta point method. Geophysical records suggest the installation of the *Nilometer* in 715 CE, a primitive device for more accurate water level measurements. As such, we selected this as the change point of significance.

For training for the Delta point method for both time series, we used a standard normal distribution prior for the Gaussian process, and hazard rate parameter *θ*_*h*_ = − 3.982. For the well-log data, the Delta point interval was chosen as 30 due to the length of the time series and sensor noise, and for the Nile river time series, the Delta point interval was chosen to be 50, as the curve is smoother. The training set consisted of the first 1000 data for the well-log series, and first 250 data for the Nile river set.

The results of each method are displayed in Table 3. The Delta point method performed better than the other methods TY, LS, and L for the well-log data set (absolute difference 2 compared to 13, 15, and 33, respectively). For the Nile river data set, all methods performed well, with declared change points of interest very close to the labelled installation of the Nilometer in 715 CE. The Delta point method for the well-log set is displayed in Figure 7 (Top Panel), and the Delta point method for the Nile river data set is displayed in Figure 7 (Bottom Panel).

**Table 3:** Well-log and Nile River results

| Method | Detected Time | Abs. Diff. |
| --- | --- | --- |
| Well-log | Labelled: 1070 |  |
| Delta | 1072 | 2 |
| TY | 1083 | 13 |
| LS | 1085 | 15 |
| L | 1103 | 33 |
| Nile river | Labelled: 715 |  |
| Delta | 720 | 5 |
| TY | 723 | 8 |
| LS | 722 | 7 |
| L | 725 | 10 |

**Figure 7:**
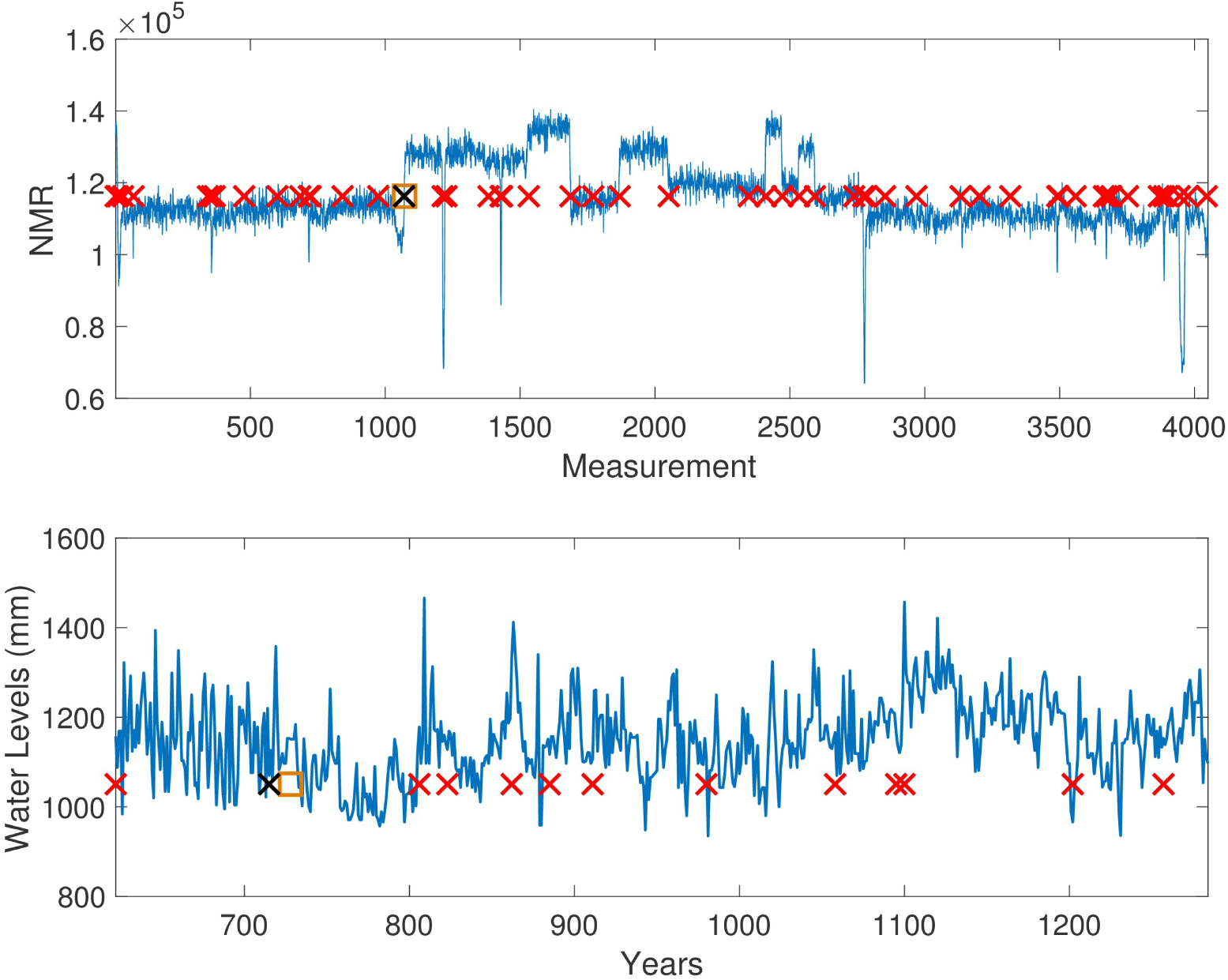
Well-log and Nile River level Delta point method. Top Panel: Delta point method for Well-log data set. The y-axis denotes NMR reading during well digging, and the x-axis denotes the measurement instance. Suspected change points are denoted with red crosses, the user-labelled change point with a black cross(1070), and the detected Delta point with an orange box(1072). Bottom Panel: Delta point method for annual lowest levels of Nile River. The y-axis denotes the water level (mm), and the x-axis denotes the years. Suspected change points are denoted with red crosses, the installation of the Nilometer(715) with a black cross, and the detected Delta point with an orange box (720).

### ECG recordings

The ECG dataset consists of short time series, with varying features. It is comprised of 100 clinical ECG recordings, each 136 data in length taken from a 67 year old patient [CKH^+^15]. Each recording contains one QRS complex for the patient, with the onset of the QRS complex user-labeled. To determine the effectiveness of the Delta point method in detecting a significant change point in short time series, we tested each method’s accuracy in detecting the QRS complex, and the difference between the labelled beginning and detected time. As each time series is short, and the QRS complex rapidly begins and ends in the recording, accurate detection of the change point was considered very important.

For training for the Delta point method, we used a standard normal distribution prior for the Gaussian process, and hazard rate parameter *θ*_*h*_ = − 3.982. Due to the short nature of these time series, the training set length was selected to be the first 30 data points; the training set never included the QRS complex for any of the 100 instances. The Delta point interval was set to 5, as the QRS complex is very short, and occurs rapidly in the series. The time series rapidly changes here, so a shorter interval performed best.

The performance of each method is displayed in Table 4. The Delta point method significantly outperformed the TY and LS method in mean absolute difference from the labeled detection times of the complex (*p* < 0.001 CI = [− 2.138,− 1.002] and [−.1339,− 0.0611], respectively). Method L had indistinguishable performance from the Delta point method (*p* = 0.954 CI = [− 0.709, 0.609]). The Delta point method had the lowest MSE of all methods (14.13 compared to 32.2, 26.31, and 22.73, respectively). A box plot of the absolute difference of all of the methods is displayed in Figure 8.

**Figure 8:**
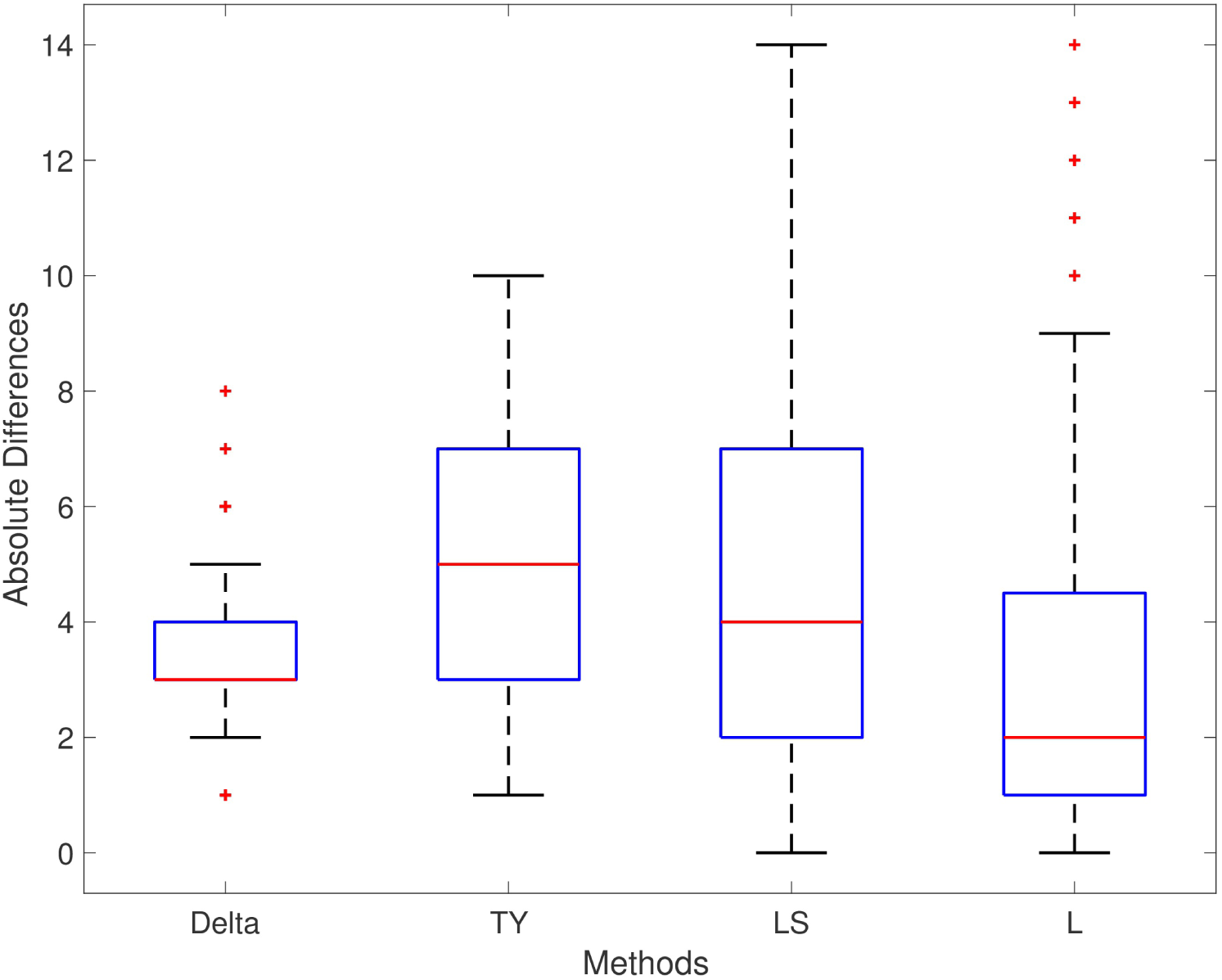
ECG (ECGFiveDays) boxplot. Legend: TY = [TY06], LS = [LS08], L = [LYCS13]. Boxplot of absolute differences of detected QRS complexes for 100 short time series of ECG recordings. The true change point is located at 0.

**Table 4:** ECG (ECGFiveDays) QRS complex results

| Method | Abs. Diff. (mean ± st. dev.) | MSE | Significance ($p$-value) [CI] |
| --- | --- | --- | --- |
| Delta | $3.51 \pm 1.352$ | 14.13 | N/A |
| TY | $5.08 \pm 2.54$ | 32.2 | 1 ($p < 0.001$) $[-2.138, -1.002]$ |
| LS | $4.21 \pm 2.944$ | 26.31 | 1 ($p < 0.001$) $[-1.339, -0.061]$ |
| L | $3.53 \pm 3.221$ | 22.73 | 0 ($p = 0.954$) $[-0.709, 0.609]$ |

Legend: TY = [TY06], LS = [LS08], L = [LYCS13]. Comparison of methods for detecting onset of QRS complex in 100 short time ECG recordings, 136 observations in length. Mean ± standard deviation of absolute difference (Abs. Diff.) between labelled times and detected time of each method are given. Mean Square Error (MSE) of each method is displayed where the lowest value displayed had the least detection error. Results of null hypothesis two-sided t-test comparing absolute differences to Delta point method displayed with *p*-values and confidence intervals [CI].

### Fetal sheep model of human labour

We applied the Delta point method to a data set consisting of 14 experimental time series of a measure of fetal heart rate variability (HRV) known as the root mean square of successive differences (RMSSD) of R-R intervals of ECG recorded during umbilical cord occlusions (UCO) [FMW^+^07, FMW^+^09]. The RMSSD may be used a measure to study the relationship between fetal systemic arterial blood pressure (ABP) and fetal heart rate in a fetal sheep model of human labour [WDR^+^14, RJA^+^13, FMM^+^09]. RMSSD is a sensitive measure of vagal modulation of HRV, and is known to increase with worsening acidemia, a dangerous condition that may occur during labour [FMW^+^07, FMW^+^09, XDR^+^14, DGB^+^14].

During UCO mimicking human labour, a hypotensive blood pressure response to the occlusions manifests as the introduction of a new trend in the recorded time series. This response is induced by the vagal nerve activation triggered by worsening acidemia during UCO as discussed in [FDG^+^15]. These points are detected by *expert* visual detection and are known as ABP *sentinel points*. These sentinel points are defined as the time between the onset of blood pressure responses and the time when pH nadir (ph *<* 7.00) is reached. A change point detection algorithm should be able to detect these sentinel points from the non-invasively obtainable fetal heart rate derived RMSSD signal in an online manner to assist in clinical decision making.

The experimental time series are short - less than 200 observations - and confounded with a large amount of noise due to experimental conditions and measurement error. The time series are piecewise locally stationary, and contain naturally occurring biological fluctuations due, for example, to non-linear brain-body interactions [BNPS97]. These factors make the detection of the expert sentinel point difficult for existing change point techniques [BP96, GP83].

To avoid false alarms, we defined a clinical *region of interest* (ROI) of 20 minutes before the sentinel point where a declared change point of interest is determined to be a success. We also took into account detections that are at most 3 minutes posterior to the sentinel point, as this is one experimental cycle late. The defined region of interest is to assist clinicians in decision making, as it provides a feasible window of time to provide clinical evaluation, as well as reject false alarm. For training for the Delta point method, we used a standard normal distribution prior for the Gaussian process, and hazard rate parameter *θ*_*h*_ = − 3.982. We trained the Delta point method with 48 data points per time series, corresponding to two hours of recording. The Delta point interval was set at 10 data, which corresponded to 25 minutes of experiment time. This interval was chosen to coincide with the clinical ROI.

The Delta point method significantly outperformed competing methods, with 11 of 14 declared change points in the ROI, compared to 3 of 14 for TY with Fisher’s exact test statistic 0.007028, 5 of 14 for LS with Fisher’s exact test statistic 0.054238, and 2 of 14 for L with Fisher’s exact test statistic 0.001838. The Delta point method applied to one animal from the data set (ID473378) is displayed in Figure 9, and the results and detection times of each method are shown in Table 5.

**Figure 9:**
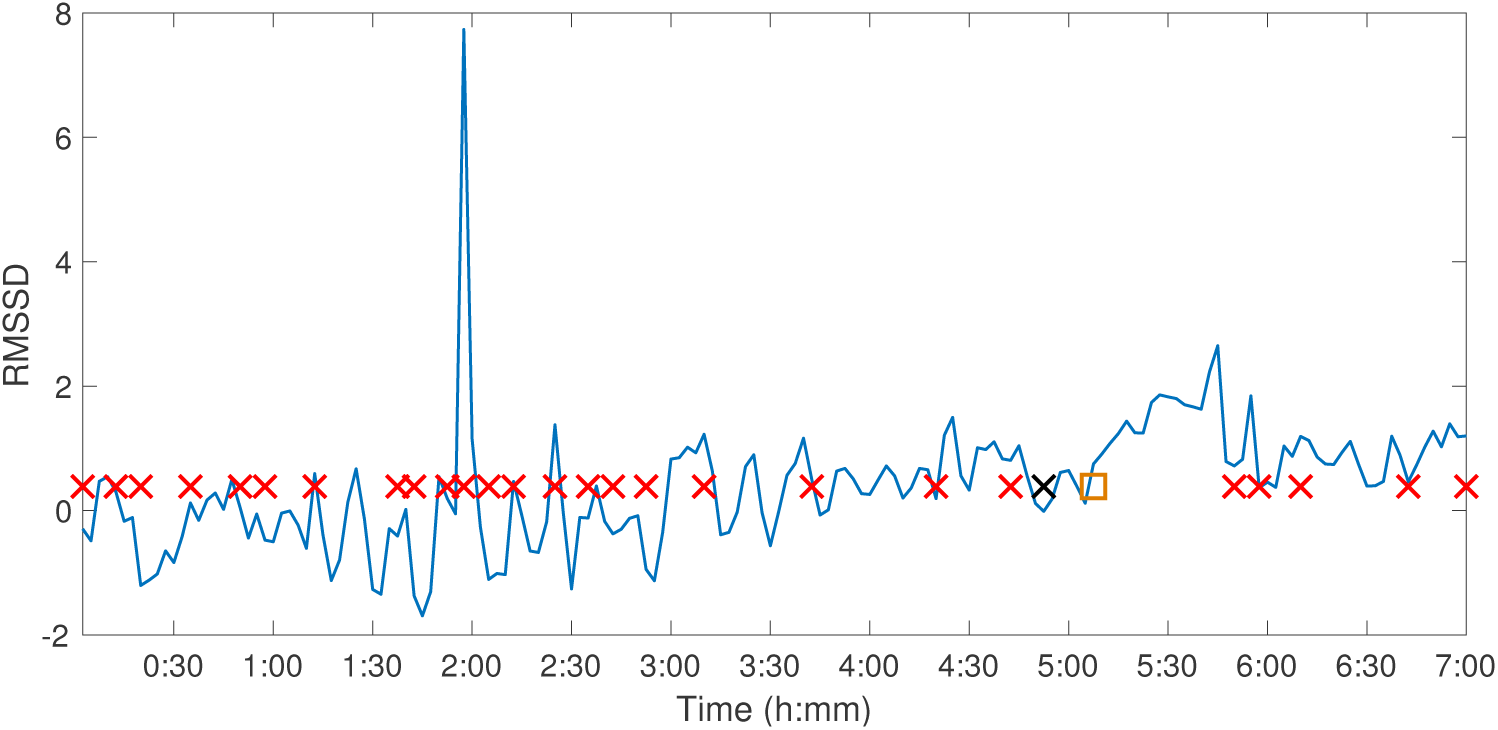
Fetal Sheep ID473378 Delta point method. Delta point method for Fetal Sheep ID473378 RMSSD time series. The y-axis denotes the RMSSD of the animal over the experimental course, and the x-axis denotes experimental time. Suspected change points are denoted with red crosses, the expert sentinel value with a black cross (6:13), and the detected Delta point with an orange box (6:24).

**Table 5:** Fetal sheep experiment results

| Animal | Sentinel(HH:MM) | Delta(HH:MM) | TY(HH:MM) | LS(HH:MM) | L(HH:MM) |
| --- | --- | --- | --- | --- | --- |
| 8003 | 15:56 | **00:07** | -00:05 | **00:12** | -00:15 |
| 473351 | 13:38 | **00:10** | -00:43 | -00:27 | 01:00 |
| 473362 | 11:05 | **00:02** | -00:48 | **-00:03** | -00:28 |
| 473376 | 12:36 | **-00:02** | **00:13** | -01:02 | -00:15 |
| 473726 | 12:04 | **00:14** | 00:25 | **00:20** | -00:10 |
| 461060 | 12:43 | **00:12** | -00:25 | 01:30 | 01:02 |
| 473361 | 12:51 | **00:15** | -00:08 | 00:35 | **00:03** |
| 473352 | 13:17 | 00:24 | 00:36 | **00:17** | -00:14 |
| 473377 | 12:12 | **-00:02** | **00:18** | **00:13** | -00:13 |
| 473378 | 13:22 | **00:13** | 00:47 | 00:37 | -00:12 |
| 473727 | 11:03 | -00:07 | **00:05** | -00:17 | -00:45 |
| 5054 | 12:53 | 01:26 | -00:30 | 01:14 | 00:44 |
| 5060 | 11:26 | **00:02** | 00:32 | 00:28 | **00:04** |
| 473360 | 13:59 | **00:07** | 01:07 | -00:17 | -00:42 |
| Total |  | 11/14 | 3/14 | 5/14 | 2/14 |

We also computed Bland-Altman plots for the experimental time series to compare the Delta point method to each other method. In Figure 10(A), we display the Bland-Altman plot for the Delta point method and TY with mean difference (6.93*±*89.03). Figure 10(B) displays the Bland-Altman plot for the Delta point method and LS, with mean difference (-1.36*±*62.6). Figure 10(C) displays the Bland-Altman plot for the Delta point method and L, with mean difference (14.4 ±59.9). In Figure 10(D), we display a modified Bland-Altman plot of the differences in detection times for each method, along with the upper and lower ROI.

**Figure 10:**
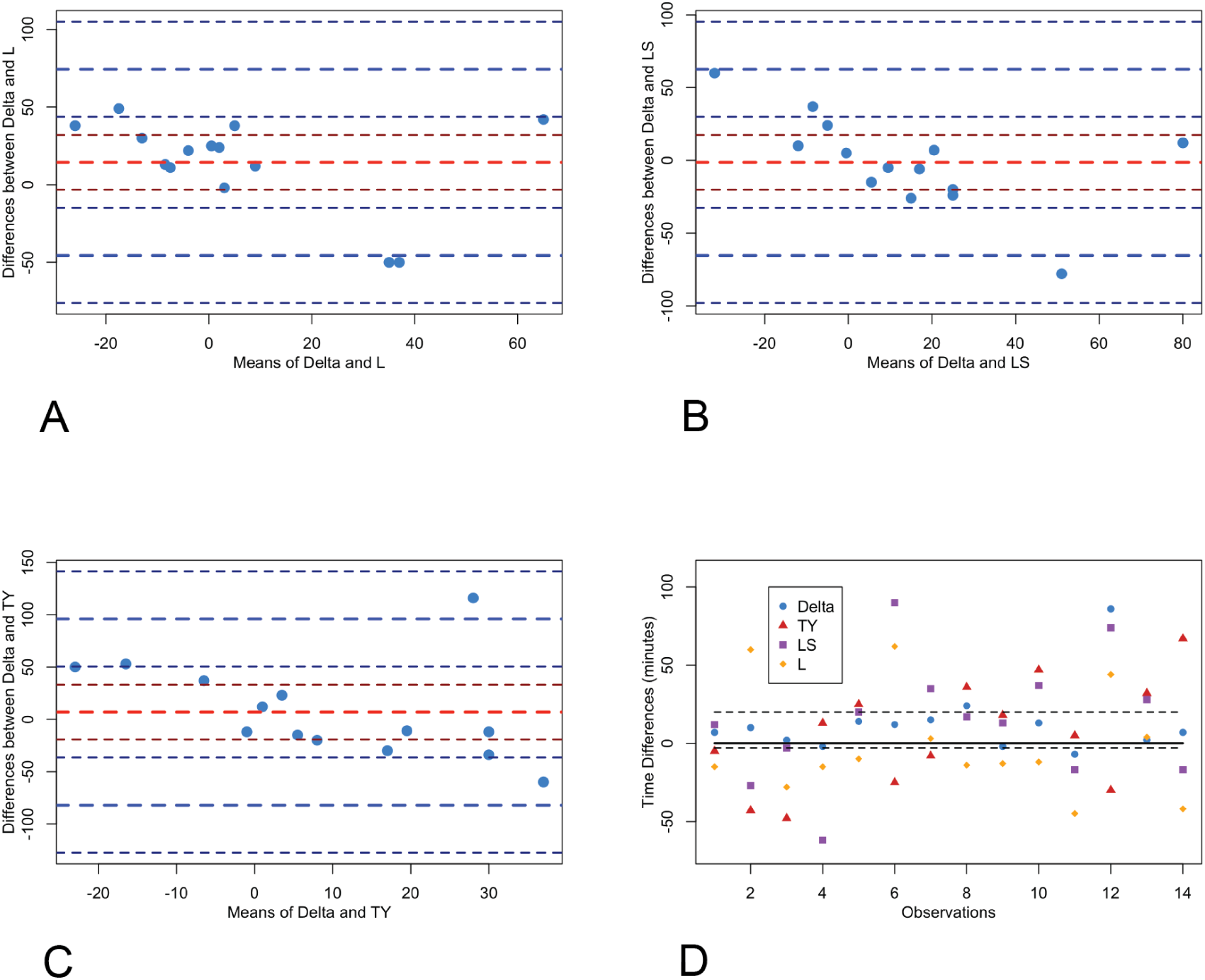
Bland-Altman plots of methods for Fetal Sheep dataset. Legend: TY = [TY06], LS = [LS08], L = [LYCS13]. Panel A-C: Bland-Altman plot comparing Delta point method to TY, LS, L, respectively. The y-axis displays the differences in the detection times, and x-axis displays the means of detection times for the two methods for each observation. The means (red) and two standard deviations (blue) are displayed with associated confidence intervals (maroon;navy). Panel D: Modified Bland-Altman plot of difference of each method and expert labelled sentinel time. Dashed black lines 20 minutes above and 3 minutes below sentinel time denote the clinical region of interest. Observations within this region are classified as a success.

## 4. Discussion

We observed that the Delta point method is effective at finding change points of interest in piece-wise locally stationary time series of different types. For the simulated time series of our own design, the Delta point method performed better or indistinguishably from the best performing methods for Series 1 and Series 2. For Series 1, the Delta point method had the lowest MSE, which suggested it is accurately identifying change points of interest. For Series 2, the Delta point method significantly outperformed the competing methods in terms of mean absolute difference in detection time for labelled change points of interest. Although method LS had a lower MSE for this series, its mean detection difference is closer to 0. In Series 3, method L performed the best, with the smallest mean absolute difference in detection time, and MSE. Series 3 consisted of the introduction of the linear trend to the autoregressive moving average model, of which the introduction of the trend was obscured by added noise. Since method L compares density ratios of the time series, its good performance on this time series is likely due to noticing these changing ratios before other methods noticed the trend.

For the Donoho-Johnstone Bump curve, the Delta point method performed nearly as well as the best performing method - method LS - with a smaller absolute difference in detection time compared to the other methods, TY and L. The Delta point method performance for the Donaho-Johnstone Block curve was better than the other methods, exemplifying the strength of the Delta point method for piecewise locally stationary time series. Our test results for the well-log data set also provides evidence of the performance of the Delta point method for piecewise locally stationary time series. For the Nile river data set, as the installation of the Nilometer is the most significant change point in the time series, and can even be noticed visually, we expected that all methods should accurately detect this change point with little variation. Indeed, our results confirm this hypothesis.

Legend: Sentinel = expert defined change point TY = [TY06], LS = [LS08], L = [LYCS13]. Comparison of methods in detecting expert defined change point. Method times are displayed relative to expert Sentinel time (HH:MM), with positive values representing change points of interest detected before the Sentinel time, and negative values representing change points of interest detected after the Sentinel time. Bolded results represent a change point of interest detected in the region of interest 20 minutes to and 3 minutes after the Sentinel time.

For the clinical ECG data set, ECGFiveDays, the Delta point method performs significantly better than methods TY and LS, however has an indistinguishable performance difference with method L, although the Delta point method has the lowest MSE. Due to the rapidly varying nature of the time series when the QRS complex begins, the ability of method L to compare density ratios between components of the time series is beneficial and improves its performance compared to other methods.

With regards to the fetal sheep experimental data set, the early detection of acidemia is better than late detection from a clinical perspective. Hence, we defined the clinical ROI according to expert physician input. The 20 minute window before the expert-labelled sentinel point provides adequate warning to clinicians to increase monitoring, or expedite delivery, while the 3 minute window posterior to the expert-labelled sentinel point is sufficiently close to be included in the experimental procedure. In clinical settings, we believe that earlier detection is better, as it provides longer decision making time, and justification for increased monitoring.

The novelty of the current work is that our method permits statistical-level predictions about concomitant changes in individual bivariate time series, simulated or physiological such as HRV measure RMSSD and ABP in an animal model of human labour. Our method is able to predict cardiovascular de-compensation by identifying ABP responses to UCO, a sensitive measure of acidosis. These predictions are reliable even in the instances when the signals are noisy. This is based on our observation that here, to mimic the online recording situation, no artefact correction for RMSSD was undertaken as is usually done for HRV offline processing [SGB11]. The two hour training time used for the Delta point method is also acceptable for delivery room settings, due to the typical time length of human labour between 6 to 8 hours on average [Alb99]. To our knowledge, no comparable statistical methods exist. Another benefit of the Delta point method is the ability to automatically extract the change point of interest with minimal user interaction, as opposed to other methods which require user specific thresholds and criteria.

Although the Delta point method performs well in settings with noise, the method is not designed to work accurately for time series that exhibit periodic structure. Due to the Gaussian process time series predictive model that is used for updating predictions, the accuracy or predictions and thus detected change points by the BOCPD algorithm depends on the kernel selected by the user. Indeed, periodic kernels do exist [EW06], however to be as general as possible, we did not implement them. Another possible limitation of the Delta point methodology is that the length of the interval for change point identification is required to be set by the user. In future work, we propose to establish an optimal window length, however this is a difficult task, as the window length should be determined both by the features of the time series, as well as the time scales over which interesting phenomena occur.

We have intentionally focused our analysis on the change point detection time, due to our interest in early detection of possibly negative phenomena in biological systems. For this reason, our analysis focuses only on the sensitivity of the method. Other methods may be more ideally suited for analysis with a certain specificity in mind. Additionally, it may be interesting to consider different time series predictive models, such as the dynamical Bayesian inference models which allow for adaptations in real-time to changes [DSMS12], or Gaussian process based Kalman filters [Tur11]. These methods provide an interesting direction for future research as they may be able to make better predictions of the time series data.

## 5. Conclusion

We have developed a novel, change point detection method for effectively isolating a change point of interest in short, noisy, non-stationary and non-periodic time series. Our method is able to effectively extract clinically relevant changes in the time series, allowing informed decision making [SGB11, See14]. By considering the joint distribution of the change points and the number of change points in disjoint intervals, the Delta point method remains robust to signal artefacts and confounding noise. We demonstrated our method on three simulated time series of our own design inspired by existing literature, curves from the Donoho-Johnstone benchmark curve data set, nuclear magnetic resonance reading from well-log measurements of geophysical drilling, annual water levels of the Nile river, a clinical ECG recording data set, and a physiological data set of fetal sheep recordings mimicking human labour. We compared the performance of the Delta point method to three existing change point detection methods. The Delta point method displays useful performance benefits in accurately extracting a meaningful change point to the user from a vector of suspected change-points.

## Additional Information

### Ethics

The human ECG dataset [CKH^+^15] was collected in agreement with the ethics procedures of the group Physical Activity Monitoring for Ageing People, https://www.pamap.org.

The animal research followed the guidelines of the Canadian Council on Animal Care and the approval by the University of Western Ontario Council on Animal Care.

### Conflict of Interest Statement

BSR and MGF are inventors of related patent applications entitled “EEG Monitor of Fetal Health” including U.S. Patent Application Serial No. 12/532.874 and CA 2681926 National Stage Entries of PCT/CA08/0058 filed March 28, 2008, with priority to US provisional patent application 60/908,587, filed March 28, 2007 (US 9,215,999). No other disclosures have been made.

### Availability of data and materials

Data and codes associated with the manuscript may be found at the following sources:

BOCPD code: https://sites.google.com/site/wwwturnercomputingcom/software/ThesisCodeAndData.zip?attredirects=0&d=1

Donoho-Johnstone data: ftp://ftp.sas.com/pub/neural/data/dojo_medium.txt

Well-log data: http://mldata.org/repository/data/viewslug/well-log/

Nile river data: http://mldata.org/repository/data/viewslug/nile-water-level/

ECGFiveDays: http://www.cs.ucr.edu/eamonn/time_series_data/

Simulation data: https://github.com/ngold5/fetal_data

## List of abbreviations used

ABP: Arterial Blood Pressure
BOCPD: Bayesian Online Changepoint Detection
ECG: Electrocardiogram
GP: Gaussian process
GPTS: Gaussian process time series
L: Liu et al changepoint detection method [LYCS13]
LS: Last and Shumway changepoint detection method [LS08]
MSE: Mean Square Error
TY: Takeuchi and Yamanishi changepoint detection method [TY06]

## Funding

This work was funded by CIHR (to MGF), FRQS (to MGF), Canada Research Chair Tier 1 in Fetal and Neonatal Health and Development (BSR); Women’s Development Council, London Health Sciences Centre (London, ON, Canada) (to BSR and MGF).

